# An improved iCLIP protocol

**DOI:** 10.1101/2021.08.27.457890

**Authors:** Flora C. Y. Lee, Anob M. Chakrabarti, Heike Hänel, Elisa Monzón-Casanova, Martina Hallegger, Cristina Militti, Federica Capraro, Christoph Sadée, Patrick Toolan-Kerr, Oscar Wilkins, Martin Turner, Julian König, Christopher R. Sibley, Jernej Ule

## Abstract

Crosslinking and Immunoprecipitation (CLIP) is a powerful technique to obtain transcriptome-wide maps of in vivo protein-RNA interactions, which are important to understand the post-transcriptional mechanisms mediated by RNA binding proteins (RBPs). Many variant CLIP protocols have been developed to improve the efficiency and convenience of cDNA library preparation. Here we describe an improved individual nucleotide resolution CLIP protocol (iiCLIP), which can be completed within 4 days from UV crosslinking to libraries for sequencing. For benchmarking, we directly compared PTBP1 iiCLIP libraries with the iCLIP2 protocol produced under standardised conditions, and with public eCLIP and iCLIP PTBP1 data. We visualised enriched motifs surrounding the identified crosslink positions and RNA maps of these crosslinks around the alternative exons regulated by PTBP1. Notably, motif enrichment was higher in iiCLIP and iCLIP2 in comparison to public eCLIP and iCLIP, and we show how this impacts the specificity of RNA maps. In conclusion, iiCLIP is technically convenient and efficient, and enables production of highly specific datasets for identifying RBP binding sites.

## Introduction

Many techniques employing high-throughput sequencing are available for ‘protein-centric’ transcriptomic studies of protein-RNA interactions, including RIP, CLIP, TRIBE, APEX-seq and others (Hafner et al. 2021). Each technique has distinct advantages as well as limitations; some define the precise position of direct protein-RNA contacts, whereas others have lower resolution but also detect RNAs that are in spatial proximity (Ramanathan, Porter, and Khavari 2019). CLIP and its derived methods fall into the first category, requiring ‘zero-distance’ covalent crosslinking of *in vivo* protein-RNA contacts, usually mediated by UVC irradiation, to enable stringent purification of the RBP-of-interest with their bound RNA (Ule et al. 2003; Lee and Ule 2018). This is coupled with library preparation and sequencing of the preserved RNA fragments.

Current variants of CLIP protocols can provide precise positional and quantitative information on the crosslink events, which is important in deciphering the regulatory functions of RBPs. These protocols vary in sensitivity and specificity due to differences in the stringency of purification to avoid re-associations during purification of protein-RNA complex, enzymatic biases, and efficiency of cDNA library preparation protocol, among others (Lee and Ule 2018). Many current CLIP variants identify crosslink positions by the analysis of cDNA truncations as established in iCLIP (König et al. 2010). Compared to the original CLIP protocol (Ule et al. 2005), iCLIP experimentally differs mainly at the adapter ligation steps. The original CLIP protocol ligated two adapters (SeqRv and SeqFw, (Lee and Ule 2018)) to the isolated RNA fragments at the 3’ and 5’ ends, respectively, whereas in iCLIP the SeqFw adapter sequence is introduced to the cDNAs during reverse transcription (RT), which establishes a way to amplify truncated cDNAs. The SeqFw adapter sequence, experimental barcode and UMIs are introduced as part of the RT primer. By amplifying the truncated cDNAs and adding the use of UMIs, iCLIP increased the coverage, nucleotide resolution and quantitative nature of the method. Since then, multiple additional ‘truncation-based’ CLIP protocols were published (Lee and Ule 2018; Van Nostrand et al. 2016; Zarnegar et al. 2016; Buchbender et al. 2019), each with its own further optimisations. This led to dozens of available variant CLIP protocols, and thus it can be challenging to select a suitable variant. Here, we present an improved iCLIP protocol, referred to as ‘iiCLIP’, where we combined several published and new optimisations to increase its sensitivity and convenience.

### Development of iiCLIP

We recently employed the iiCLIP protocol for the comparative studies of RBP binding upon changes in condensation. The reproducibility of this streamlined approach allowed us to compare across a panel of TDP-43 C-terminal mutant variants, and thereby derive principles for condensation-dependent assembly on RNA-binding regions which have unusually long stretches of dispersed UG-motifs (Hallegger et al. 2021). The rationale of the improvements compared to iCLIP are explained in Table 1; the schematic overview of the full protocol is found in Figure 1. Notably, iiCLIP offers an alternative to radioactive labelling of RNPs, which was first introduced by irCLIP for quality control at the SDS-PAGE step (Zarnegar et al. 2016), while retaining similar enzymatic steps of iCLIP but with improved efficiency, combined with Ampure beads-based purifications to streamline the library preparation procedure and minimise loss of material. Thus, the resulting data from iiCLIP experiments is expected to have similar or better resolution and specificity compared to iCLIP, with greater efficiency in library preparation, and we confirmed this in the benchmarking comparisons presented below (Figures 2-4).

**Table 1:**
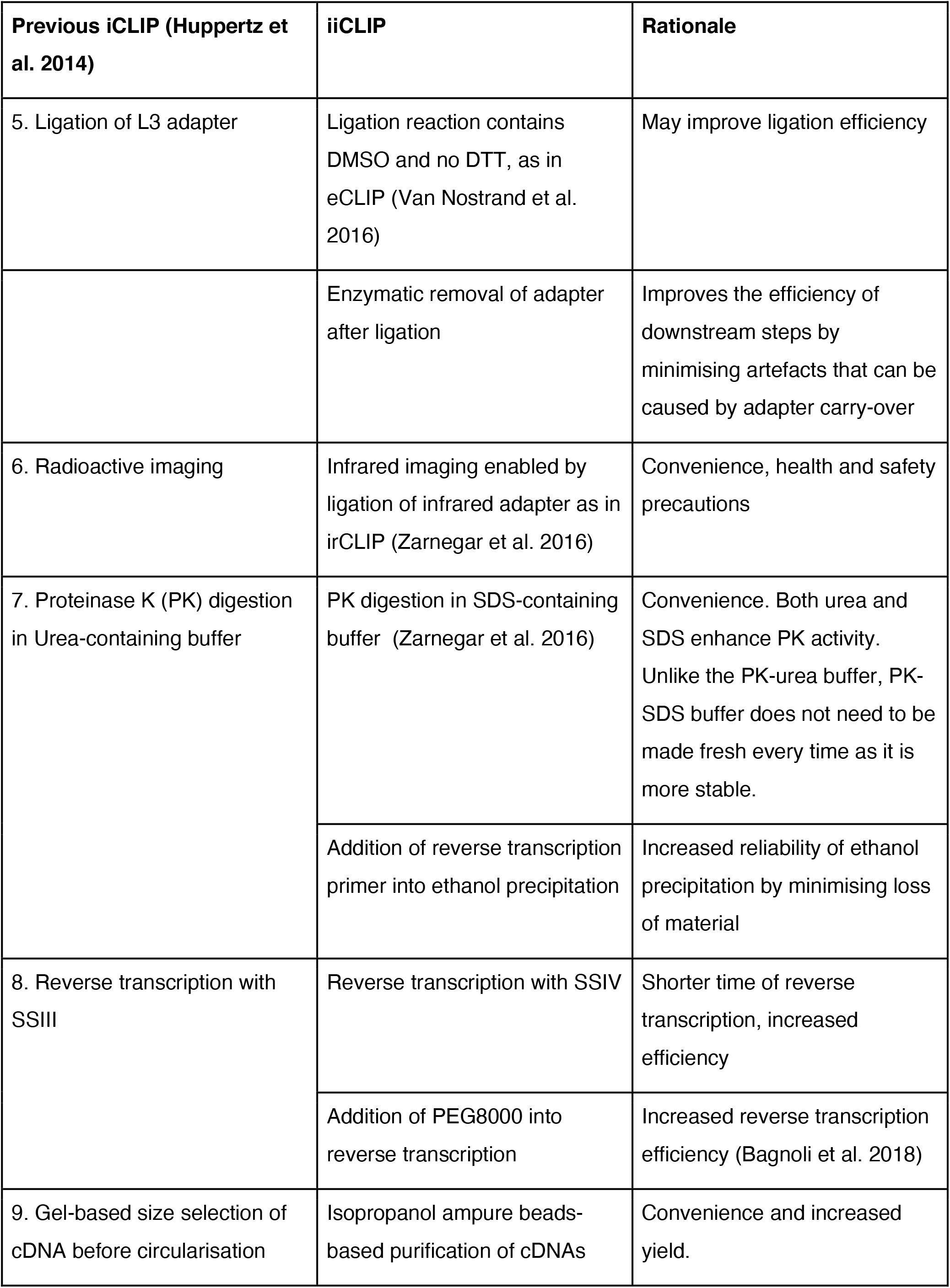
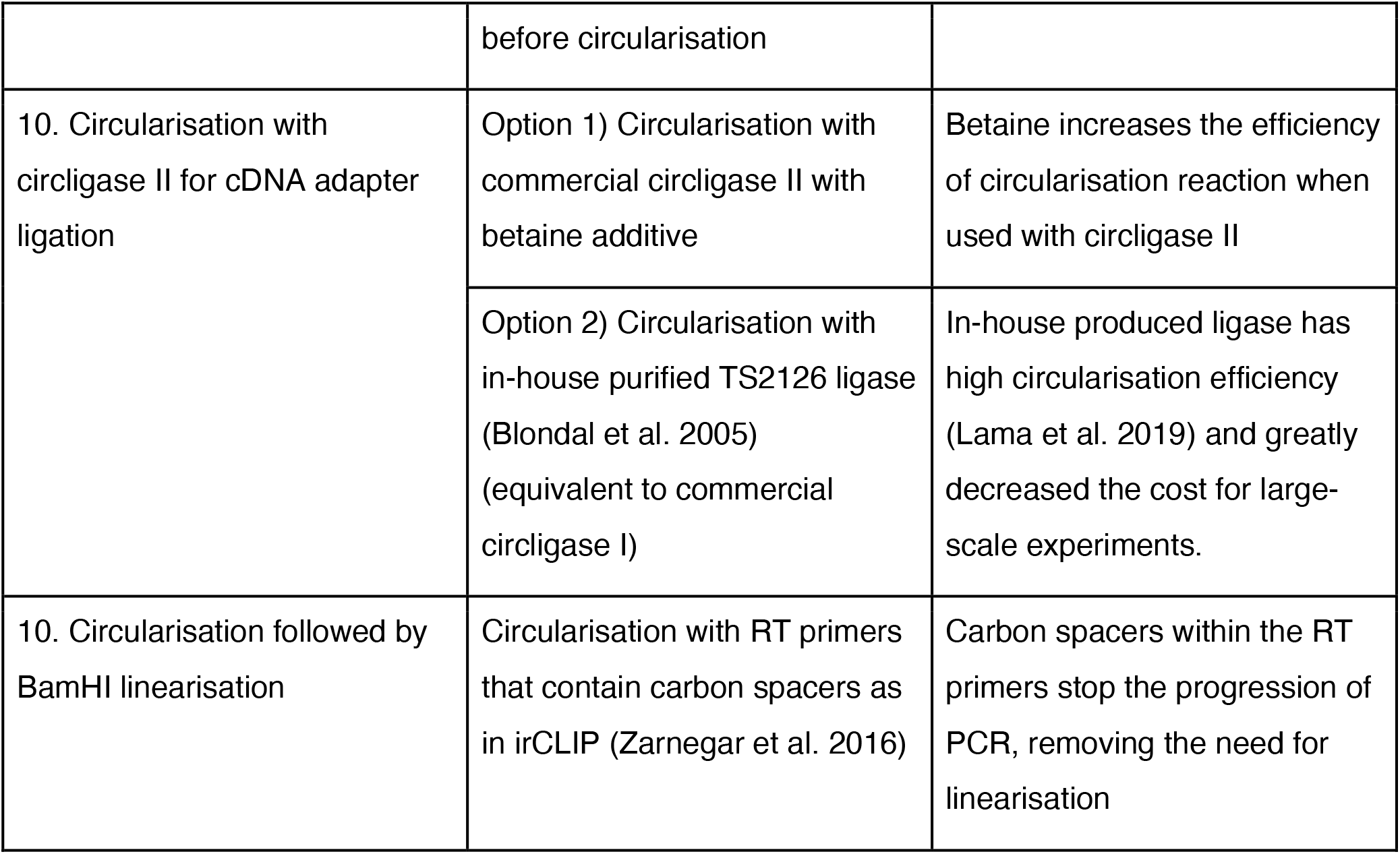
summary and rationale of optimisations

**Figure 1:**
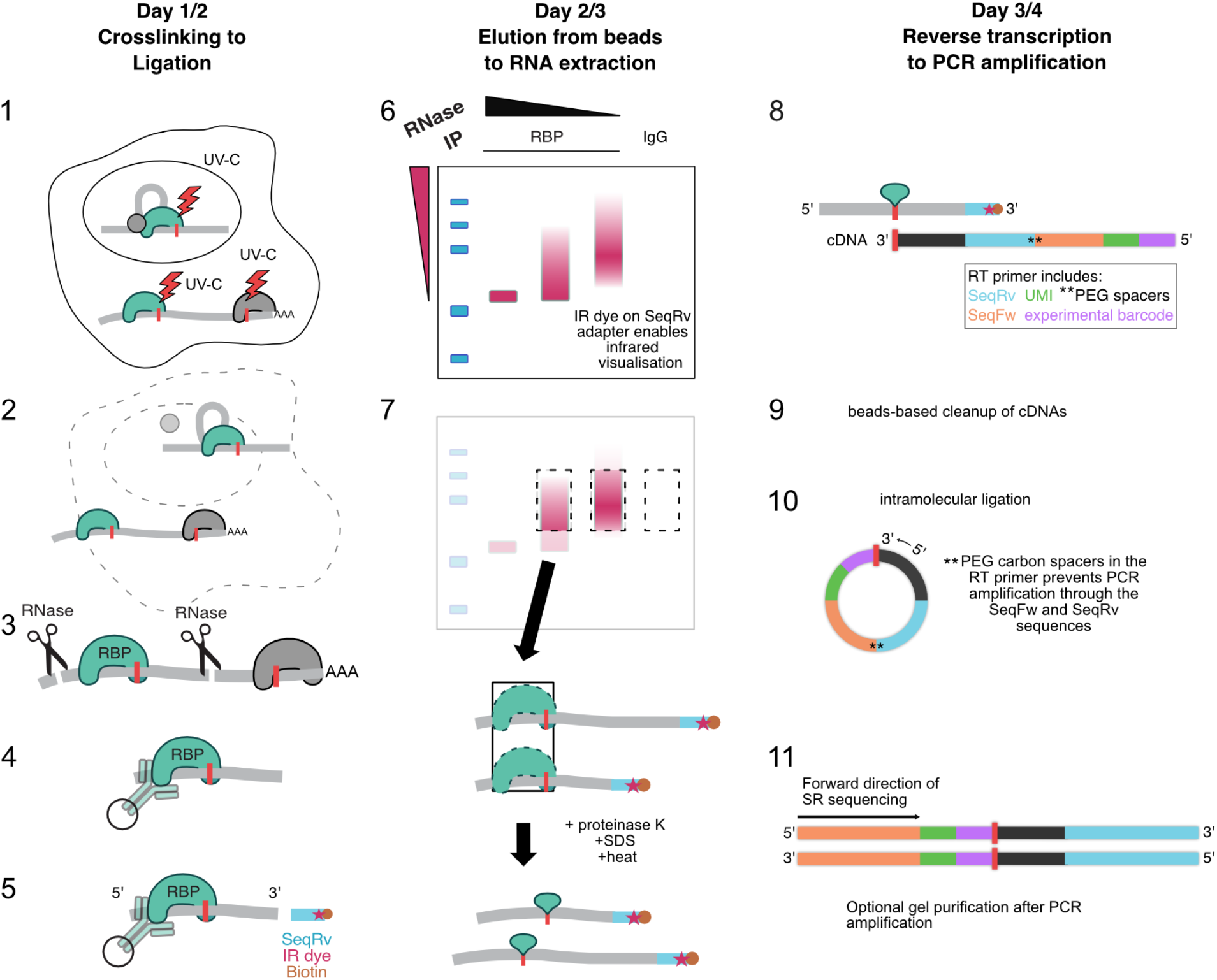
iiCLIP overview. Schematic of iiCLIP. There are 11 main experimental stages in CLIP and variant protocols, as previously defined in Lee and Ule (2018). These include: 1. Covalent protein-RNA crosslinking 2. Cell lysis 3. RNA fragmentation 4. Purification of protein-RNA complexes 5. Ligation of SeqRv adapter to fragmented RNA 6. Quality control by visualisation of captured protein-RNA complexes 7. RNA extraction 8. Reverse transcription 9. cDNA purification 10. SeqFw adapter ligation 11. cDNA amplification followed by high-throughput sequencing of multiplexed libraries. Stages where key changes have been introduced are described in Table 1. The protocol can be completed from UV crosslinking to library preparation in 3/4 days.

**Figure 2:**
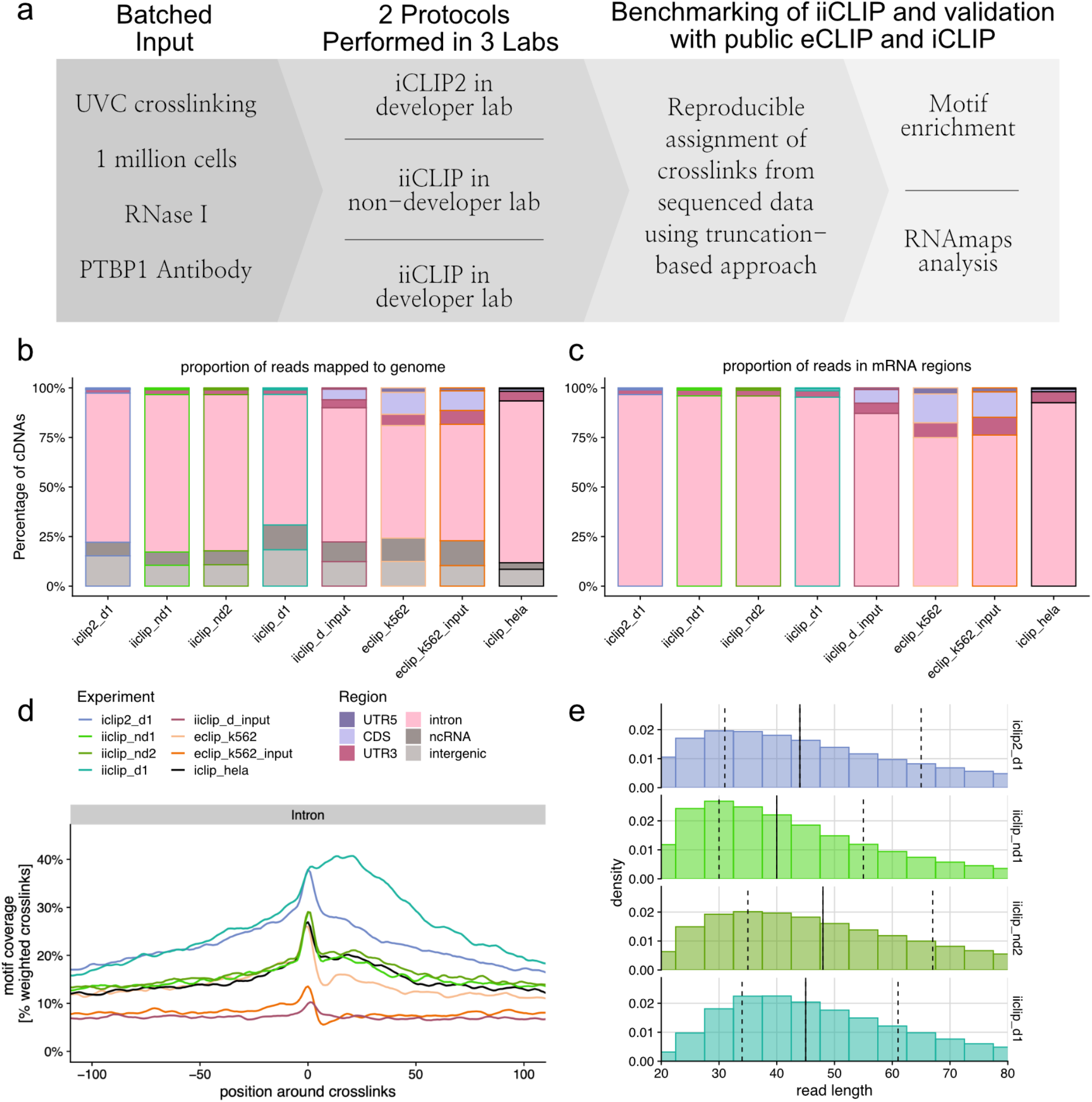
Regional and motif enrichment in identified raw PTBP1 crosslink positions. a) Design of iiCLIP benchmarking comparisons. Input material is controlled appropriately between iiCLIP and iCLIP2 libraries, and downstream bioinformatic analysis is done systematically, including for public eCLIP and iCLIP data. (b-c) Regional distribution of cDNAs mapped to (b) genome or (c) filtered for mRNA regions (CDS, 5’UTR, 3’UTR, intron) identified by the different protocols and experiments. d) Enrichment of known PTBP1 pentamers around crosslink positions (% crosslinks weighted by cDNA counts with motif coverage) in intronic regions. (e) Histogram showing read length distribution of uniquely genome-mapping reads. Solid line indicates the median read length, and dashed lines indicate the lower and upper quartiles.

**Figure 3:**
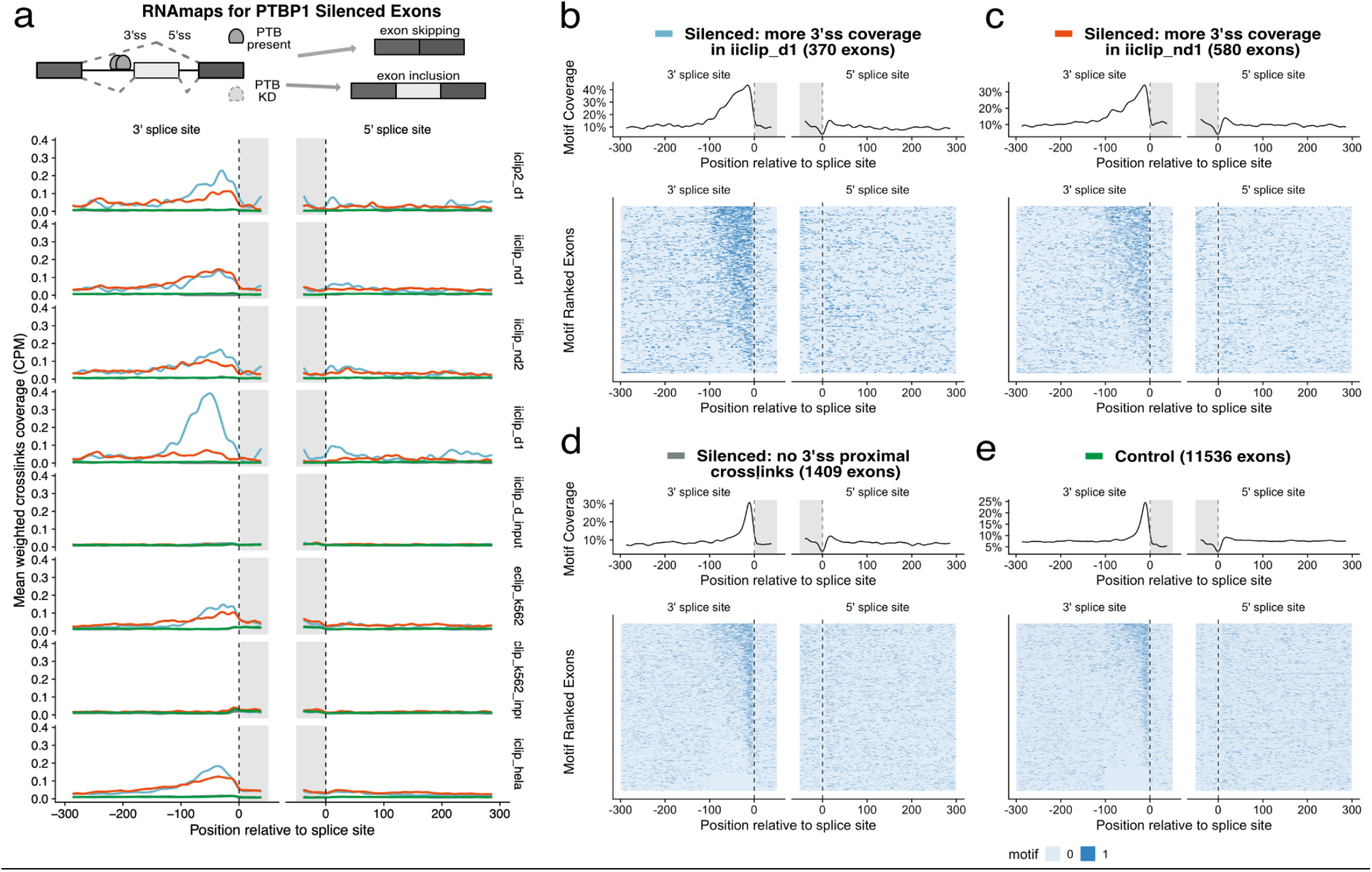
PTBP1 RNA maps evaluate the sensitivity and specificity across methods. a) In the following PTBP1 RNA maps, weighted crosslinks normalised by library size are plotted as a metaprofile around the 3’ and 5’ splice sites of PTBP1 silenced compared with control (Green) exons identified from public PTBP1/2 KD RNAseq. Silenced exons were stratified into 3 classes based on crosslink site coverage proximal to the 3’ss (−100..0 window). Light blue: more covered in iiclip_d1 compared to iiclip_nd1; Orange: more covered in iiclip_nd1 compared to iiclip_d1; Grey: no detected crosslinks from both libraries. The profile is used to evaluate the specificity and sensitivity of each method. b-e) Motif maps for each of the 3 classes of PTBP1 silenced exons (b-d) and the control exons (e), shown as a coverage metaprofile (top) or heatmap (bottom). (b) More covered in iiclip_d1 compared to iiclip_nd1; (c) more covered in iiclip_nd1 compared to iiclip_d1; (d) no detected crosslinks from both libraries.

**Figure 4:**
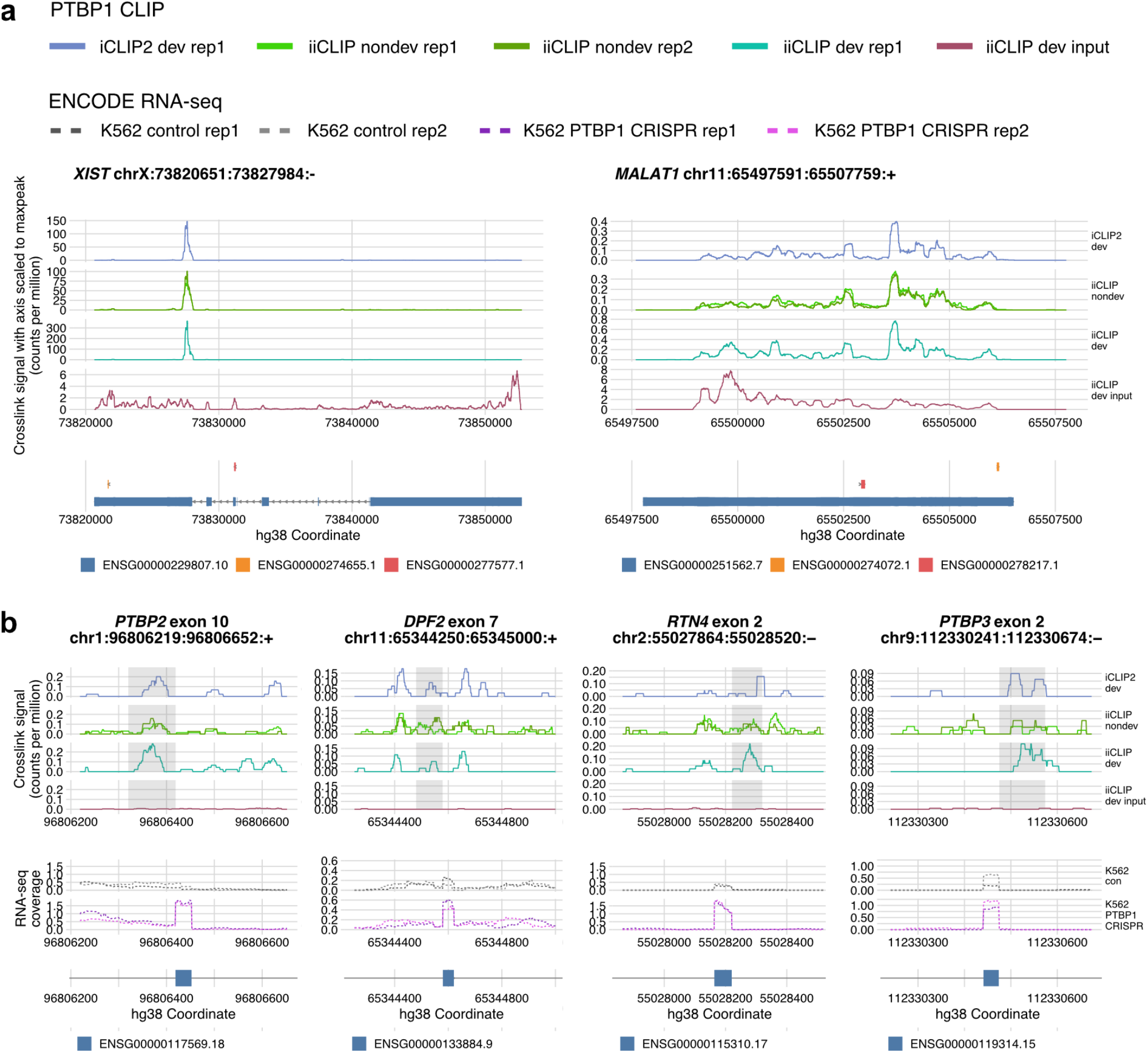
Visualisation of PTBP1 iCLIP2 and iiCLIP crosslink events on PTBP1 RNA binding sites. (a) Comparative visualisation of PTBP1 CLIP libraries on ncRNA targets *XIST* and *MALAT1*. Top panel: library size normalised crosslink counts from PTBP1 CLIP libraries generated for this study are plotted. Axis is scaled by the maximum peak height of each group to visualise the distribution of crosslink signal in the input library, containing global protein-RNA crosslinking signal, compared to PTBP1 crosslink signal. Bottom panel: Gene annotation (GENCODE hg38 v27). (b) Comparative visualisation of PTBP1 CLIP libraries on individual intronic sites surrounding regulated exons. Top panel: library size normalised crosslinking counts from PTBP1 CLIP libraries generated for this study are plotted, centred on examples of previously validated PTBP1 repressed cassette exons. The region of -100…0 upstream of the 3’ splice site is shaded in grey, as this region contains the highest reproducible enrichment of crosslinks across silenced exons as shown previously in the RNAmaps metaprofiles. Middle panel: RNAseq coverage (CPM) from ENCODE K562 control and PTBP1 CRISPR libraries. Cassette exon inclusion increases upon PTBP1 CRISPR treatment. Bottom panel: Gene annotation (GENCODE hg38 v27).

### Benchmarking of iiCLIP: a controlled comparison using the same input material

To assess the performance of iiCLIP, we performed several comparisons. First, to test that iiCLIP can be easily adopted by additional labs, we designed a comparison between CLIP protocol developer and non-developer labs. Two iiCLIP libraries were generated in a non-developer lab, and a single iiCLIP library was produced in the iiCLIP developer lab for the purpose of this study (Supplementary Table 1). Second, we directly benchmarked iiCLIP against the recently published iCLIP2 protocol (Buchbender et al. 2019). iCLIP2 was shown to increase the sensitivity of iCLIP by an order of magnitude and can be completed within four days (Buchbender et al. 2019). Like iCLIP, iCLIP2 contains the radioactive labeling step to monitor optimal conditions of immunoprecipitation and RNA fragmentation. The iiCLIP protocol differs from iCLIP2 in the non-radioactive visualisation of the protein-RNA complex, conditions of RNA ligation, cDNA ligation and in the post-RT conditions of cDNA purification and amplification.

These comparisons were controlled for using the same input cells, antibodies and RNase treatment. A batch of cells was prepared and UVC crosslinked in the Ule lab, which were sent along with a tested batch of RNase and antibody to two other laboratories. Thus, the experiments were performed with identical RNase treatment and immunoprecipitation conditions. After sequencing using the described conditions (Table S1), reads were processed through the primary analysis pipeline on the iMaps web server (imaps.genialis.com/) to identify crosslink positions by mapping cDNA truncations and to quantify cDNA counts per position using UMIs, which applies for all protocols compared in this study (Figure2A, Tables S2 and S3).

### Validation of sensitivity and specificity of data through PTBP1 motif analysis

The CLIP method has been developed with the purpose of obtaining data with high specificity for the direct RNA binding sites of an RBP. An additional aim during iiCLIP development has been to maximise data sensitivity by increasing the complexity of the produced cDNA library, and thus its capacity to comprehensively identify binding sites across a wider dynamic range of RNA expression and with less input material. However, it is important that changes in data specificity are carefully assessed whenever changes are made that aim at higher sensitivity (Hafner et al. 2021). Here, we wished to gain a comparative perspective on both the sensitivity and specificity of data obtained by both protocols, which we produce in this study, and with representative public data from other published CLIP variants. We focused on PTBP1, a well-characterised nuclear RBP and splicing regulator that has been used by several labs to benchmark their own CLIP variants, which has already enabled some meta-analyses based on these published data (Chakrabarti et al. 2018; Haberman et al. 2017; Van Nostrand et al. 2016; Zarnegar et al. 2016; Gu et al. 2018). Specificity of PTBP1 CLIP data can be assessed through sequence motif enrichment around crosslinks, as its binding to UC-rich RNA motifs has been well characterised with *in vitro* biochemistry and structural studies, and is supported by evidence of motif enrichment around regulated exons (Oberstrass et al. 2005; Spellman et al. 2005; Chou et al. 2000; Cereda et al. 2014). Sensitivity for functionally important sites can be shown with an RNA map (Ule et al. 2006), which demonstrates how PTBP1 binding around regulated exons has bidirectional outcomes on RNA processing in a position-dependent manner (Witten and Ule 2011; Xue et al. 2009; Llorian et al. 2010). Therefore the analysis of PTBP1 CLIP data can provide information on sensitivity and specificity of the crosslink positions and binding peaks through multiple means (Chakrabarti et al. 2018), which we have exploited here for comparative analysis of CLIP variants.

We produced iCLIP2 and iiCLIP data of PTBP1 immunoprecipitation from 1 million HEK293 cells for the purpose of controlled comparison. For the purpose of nomenclature, we refer to data produced by developer labs as iCLIP2_d and iiCLIP_d, whereas the data produced in non-developer lab is referred to as iiCLIP_nd. To contextualise these new data with existing protocols, we reprocessed public iCLIP and ENCODE eCLIP raw data on PTBP1 with the identical computational pipeline on iMaps (Table S3). First, we examined the regional distribution of sequenced cDNAs (Figure2B, C). As expected, iCLIP2 and iiCLIP PTBP1 cDNAs predominantly map to introns, similar to public iCLIP (Figure 2B, C). Interestingly, we observed differences in the proportion of reads mapping to intergenic and non-coding RNAs, which is highest in iiCLIP_d1 (18% and 12%) and lowest in public iCLIP (3% and 8%) (Figure 2B). On the other hand, public eCLIP data has a ∼10-fold greater proportion of CDS reads (11% in eCLIP compared to 0.3-1.2% in other variant protocols). Therefore, whilst the regional distribution of sequenced reads can be protocol- and experiment-dependent, iCLIP2, iiCLIP and public iCLIP are generally similar when comparing cDNAs which map to mRNA regions (Figure 2C).

Next, we assessed the occurrence of PTBP1-binding motifs around the intronic crosslink positions. As is standard for iCLIP and eCLIP, crosslink sites were assigned to the genomic coordinate preceding the start of cDNA inserts. We then evaluated the coverage around crosslink positions of 12 UC-rich pentamers that were found to be most enriched in previous PTBP1 CLIP studies (Haberman et al. 2017). These pentamers were highly enriched at the crosslink positions of all datasets evaluated here, but with major differences in the extent of enrichment in the region within 10nt of crosslink sites, from the minimum of 18.7% (in eCLIP) to the maximum of 37.9% (in iiCLIP_d) (Figures 2D and S1E). We were surprised to find that the iiCLIP_nd had lower enrichment than iiCLIP_d, such that iiCLIP_nd was similar to the public iCLIP (19.3-20.6%). Reassuringly, motif enrichment around 3’UTR, intergenic and ncRNA crosslink positions showed similar variability across the libraries as in introns, and the strong occurence of PTBP1 motifs generally supported the specificity of crosslink sites identified in all of these regions (Figure S1A-C, E). However, eCLIP crosslinks in CDS did not show motif enrichment compared with SMInput data, suggesting that the increased abundance of CDS cDNAs may derive from background contamination (Figure S1D). Of additional interest, the highest motif enrichment of all regions was seen in the intergenic RNA crosslinking sites (Figure S1A), which were most represented in iiCLIP_d, highlighting the need to better understand the sources and functionality of these sites (Agostini et al. 2021).

Notably, we also observed major variations in the broader profiles of motif enrichment around crosslink sites. In eCLIP, motif enrichment was higher upstream than downstream of crosslink sites, while in all other datasets the enrichment was higher downstream of crosslink sites (Figures 2C and S1). We believe this is likely the result of higher crosslinking energy used in eCLIP (400mJ/cm^2^), as compared with all other variants (150mJ/cm^2^) (Van Nostrand et al. 2016; Huppertz et al. 2014). Namely, high crosslinking increases the chance for multiple PTBP1 crosslink events on binding regions that contain a sequence of multiple motifs, and as a result, cDNAs will truncate closer to the 3’end of the bound RNA regions. Moreover, the iiCLIP_d and iCLIP2 datasets have the broadest enrichment of motifs. Specifically, in the region 11-50nt downstream of crosslink sites, average motif enrichment is 14.3% (eCLIP), 17.5-19.5% (iCLIP and iiCLIP_nd), 23.7% (iCLIP2) and 36.5% (iiCLIP_d). It has been shown that such broad patterns of enrichment can be derived from highly multivalent, long binding regions, which are more challenging to map in the genome due to their repetitive nature, and therefore long cDNAs are required to detect crosslink sites in these regions (Haberman et al. 2017). Hence, the shorter read length of cDNAs in iiCLIP_nd1 may contribute partly to differences in the ability to identify crosslinks in the highly multivalent UC-rich regions (Figure 2E).

### Examination of data specificity and sensitivity by RNA maps analysis

As a further independent approach to assess the specificity and sensitivity of community CLIP experiments, we examined their capacity to identify *in vivo* RNA binding that is relevant to the position-dependent regulatory role played by PTBP1 in alternative splicing. For this, we analysed RNA maps of PTBP1 for the different CLIP variants, which examines RBP binding sites around the exons that are differentially spliced upon depletion of the RBP (Figures 3 and S2) (Ule et al. 2006). We used exons which were identified either as silenced or enhanced by PTBP1 from RNA-seq analysis of HEK293 cells after PTBP1/2 knockdown (Gueroussov et al. 2015) (Figures S2A,3), microarray analysis upon PTBP1/PTBP2 knockdown in HeLa cells (Llorian et al. 2010) (Figure S2B), or from ENCODE RNA-seq analysis of K562 cells after PTBP1-targeted CRISPR treatment (Van Nostrand et al. 2020) (Figure S2C). We analysed the coverage of crosslink events up to 50nt into the exonic and 300nt into the intronic sequence flanking the splice sites of the affected exons. The strongest enrichment of PTBP1 crosslink events was seen in the upstream -100…0 window of the 3’ splice site of PTBP1-silenced exons, as compared to control exons (Figure S2A-C). This enrichment could be observed regardless of whether the PTBP1 regulated exons were defined from analysis of cell-line matched datasets (HEK293 for iiCLIP and iCLIP2 libraries, HeLa for public iCLIP and K562 for public eCLIP data).

To further understand how different datasets perform in detecting the proximal 3’ splice site enrichment pattern around silenced exons, we classified silenced exons from the HEK293 RNA-seq based on the relative coverage of crosslinks from the iiCLIP_d1 and iiCLIP_nd1 dataset in the 100nt window upstream of the 3’ splice site and analysed the RNA maps crosslink metaprofiles for these classes separately as a measure for sensitivity (Figure 3A). Silenced exons which were detected preferentially in iiCLIP_d1 had slightly broader (up to -100 relative to 3’ splice site) and higher PTBP1 motif coverage compared to those exons with relatively greater crosslink coverage in iiCLIP_nd1 (Figure 3B&C), which likely reflects the differences in motif coverage around crosslink events in the two datasets (Figure 2C). Notably, a third class of PTBP1-silenced exons that had no detectable crosslinks in both datasets showed a lower enrichment of motifs restricted to a sharp peak directly upstream of the 3’ splice site, a pattern of motif coverage more similar to the control exons (Figure 3D&E). The difference in crosslinking coverage for the three classes of silenced exons is reproduced across all PTBP1 datasets (Figure 3A). Thus all CLIP variants preferentially identified silenced exons harbouring multivalent RNA-binding regions upstream of the 3’ splice site, which are candidate exons for direct PTBP1 repression.

Finally, we compared the PTBP1 crosslinking profiles identified by CLIP variants at individual RNAs. All iiCLIP and iCLIP2 libraries produced for this study show a highly reproducible distribution of crosslink positions and frequency on the ncRNAs XIST and MALAT1 (Figure 4A). Compared to the input control produced in parallel to iiCLIP (iiclip_d_input, see methods), all of these PTBP1 libraries show ∼hundred-fold enrichment of signal in the E repeat region of Xist, whereas the normalised signal across almost all positions of MALAT1 was lower in iiCLIP compared to input control, in agreement with the known functional binding of PTBP1 to Xist (Pandya-Jones et al. 2020) but not MALAT1. We next analysed the RNAs containing PTBP1-repressed exons, where we also observed highly reproducible crosslink signal across all the CLIP libraries produced in this study (Figure 4B). Thus, by performing comparative CLIP analysis across multiple labs using controlled batches of input material and immunoprecipitation conditions, we show that both protocols are capable of generating highly specific data with similar sensitivity in spite of their major downstream differences in the conditions of reverse transcription, cDNA ligation enzymes, strategies of cDNA purification and PCR amplification. In summary, the iiCLIP protocol can be set up and completed effectively in a non-developer lab, and overall shows similar sensitivity and specificity to the latest variant CLIP protocols.

## Discussion

### Sources of biological and technical background in CLIP

A critical aspect of CLIP data analysis is to derive confident binding sites from the crosslink events detected in the experiment, by identifying positions with signal above the background distribution (Chakrabarti et al. 2018). There are two sources of background to be considered: intrinsic and extrinsic. Biological noise in CLIP data can arise from true RBP crosslinking capturing stochastic RBP-RNA interactions which occur in addition to binding at cognate high-affinity sites, which can be referred to as ‘intrinsic background’ (Hafner et al. 2021). Extrinsic background, however, partly derives from RNA fragments crosslinked to co-purified RBPs. Such background can be relevant for understanding the function of the RBP-of-interest in case the co-purified RBPs are interacting partners within larger RNP complexes, though it may obscure the intrinsic specificity of the RBP. Another type of extrinsic background derives from sources which do not biologically relate to the RBP-of-interest, for example contamination with abundant RNA fragments or abundant cellular RBPs with their crosslinked RNAs (Hafner et al. 2021). This is most likely to be cell-state and protocol-dependent. In addition to these sources of background, CLIP data are affected by technical biases in the library preparation protocol. These dynamically alter the distribution of final sequenced reads such that their observed abundance in a CLIP experiment may not faithfully represent the repertoire of isolated RNA fragments.

One approach to assess the extrinsic background of CLIP is to sequence input libraries, as done routinely for eCLIP SMInput samples that loads the crosslinked lysate on the gel and isolates RNA from a size-matched region of the membrane followed by in-solution dephosphorylation and adapter ligation. Here we used a modified approach to producing input data, where crosslinked protein-RNA complexes are captured first on beads (Moggridge et al. 2018), allowing parallel processing of both IP and input samples and on-bead dephosphorylation and adapter ligation, which makes the conditions of the input sample fully comparable to the IP sample in the iiCLIP protocol. We procured iiCLIP_d_input in conjunction with the PTBP1 iiCLIP_d1. Similar to the eCLIP SMInput, the iiCLIP_d_input sample also shows some enrichment in CDS exons (Figures S2A, 1B). However, we observed negligible overlap in the signal between the PTBP1 iiCLIP_d and corresponding input libraries, both at the metaprofile RNA splicing map level and individual binding site level (Figures 3, 4, S2). While the CDS enrichment from PTBP1 eCLIP warrants background normalisation by eCLIP SMInput, there is no CDS crosslink enrichment in PTBP1 iiCLIP and thus input normalisation is not required to remove such signal. Instead, peak-calling approaches that model the intrinsic background from iiCLIP data itself might be more appropriate for iiCLIP, and input data could instead serve as a proxy to account for RNA abundance (Hafner et al. 2021).

### Limitations of iiCLIP protocol and future directions

We showed iiCLIP has good sensitivity for 1 million cells, however when input material is low, adapter-primer and RT primer concatemer artefacts do emerge during PCR in iiCLIP libraries. This indicates that whilst we have introduced enzymatic removal steps after ligation and reverse transcription, they are insufficient to remove all excess oligos. In cases of visible artefact contamination, these artefacts should be removed by gel purification after PCR amplification, which minimises the loss of unique cDNAs, prior to sequencing. Further optimisations on adapter and RT primer sequences and/or annealing temperatures might minimise these artefacts.

Moreover, iiCLIP shares similar limitations with other CLIP variant protocols (Lee and Ule 2018). The CLIP approach relies on a specific antibody. If this is not available, it may be possible to express epitope-tagged proteins in the relevant cells lines or other types of input material. Orthogonal methods to CLIP may also be applicable, such as ones which rely on mapping of RNA-tagging or editing events (Lapointe et al. 2015; McMahon et al. 2016; Brannan et al. 2021). These are especially valuable if the protein of interest does not crosslink well, for example when indirect protein-RNA interactions are of interest to study functions of a protein-complex. CLIP can also be used comparatively to understand selective remodelling of RNP networks in *vivo (Hallegger et al. 2021)*, and to take full advantage of such analyses, further development of experimental and computational approaches will be valuable.

As CLIP protocols continue to evolve, it is important to assess both sensitivity and library specificity with computational strategies using a benchmarking RBP or ideally multiple RBPs with different binding characteristics. Accordingly, we advocate caution in the interpretation of improvements which are primarily based on measurements of library depth. Resolving the next experimental and computational challenges for CLIP technologies will allow us to design robust experiments with limited input material to probe dynamic biological contexts, thereby advancing our understanding of the principles underlying the assembly of protein-RNA complexes and their coordinated regulatory functions.

## Author Contributions

F.C.Y.L. and J.U. conceptualised the project. F.C.Y.L., M.H., C.M., F.C., C.Sa., P.T.-K., O.W., C.R.S. contributed to iiCLIP protocol optimisations. J.U. supervised iiCLIP development. F.C.Y.L. prepared CLIP materials for benchmarking. F.C.Y.L., H.H.,E.M.-C. performed CLIP experiments for benchmarking. A.M.C. provided software. A.M.C. and F.C.Y.L. performed CLIP benchmarking analysis. J.U. supervised CLIP analysis. M.T., J.K. and J.U. supervised CLIP benchmarking experiments. F.C.Y.L. and J.U. wrote the paper and protocol with input from all other authors.

## Acknowledgements

The authors thank Charlotte Capitanchik for providing the HEK293 RNAseq alternative splicing analysis used for benchmarking and Klara Kuret for help on motif coverage; Jan Medenbach for providing the TS2126 RNA Ligase I (Circligase) expression plasmid; Ina Huppertz, Tom Schultz, Matthias Hentze, and the past and current members of Ule lab for valuable discussions and assistance. This research was funded in whole, or in part, by the Wellcome Trust [103760/Z/14/Z, 215593/Z/19/Z (to J.U.); 105202/Z/14/Z (to F.C.Y.L.)]. For the purpose of open access, the author has applied a CC BY public copyright licence to any Author Accepted Manuscript version arising from this submission. This research was funded additionally by the ERC [617837-Translate and 835300-RNPdynamics (to J.U.)]; and Biotechnology and Biological Sciences Research Council (BBSRC) grants [BB/P01898X/1 and BBS/E/B/000C0428 (to E.M-C. and M.T.)]. The Francis Crick Institute receives its core funding from Cancer Research UK (FC001110), the UK Medical Research Council (FC001110), and the Wellcome Trust (FC001110).

## Methods

### Data availability

Benchmarking datasets in this manuscript are described in Table S1 and S2. Sequencing data newly generated has been deposited to ArrayExpress (E-MTAB-10881). Processed CLIP data can be accessed on the iMaps webserver (https://imaps.goodwright.com/collections/1203/).

### Code availability

Code used to generate all figures for iiCLIP benchmarking can be found on the github repository of this manuscript (https://github.com/ulelab/iiclip-benchmarking).

*Shared conditions, cell pellets and reagents across benchmarking CLIP experiments against PTBP1* HEK293 cells were grown in monolayer to 90% confluency on 150mm dishes with DMEM 10% FBS media prior to UVC crosslinking. Cells were washed once with 10ml of 1x PBS, and UVC crosslinked (254nm) at 150mJ/cm^2^ on an ice-filled tray with a Stratalinker 2400. Cells are then scraped off the dish on ice in PBS, and spun down at 250G at 4°C. Subsequently, cells are resuspended in 10ml of 1x PBS and counted with Countess automated cell counter (ThermoFisher). 1 million cells were distributed to individual eppendorf tubes, pelleted at 500G at 4°C, and snap frozen on dry ice to be stored at -80°C. For distribution, cell pellets and RNase I (EN0602, Thermofisher scientific) aliquots were sent on dry ice, whereas mouse monoclonal anti-PTBP1 antibody (sc-56701, santacruz) were sent on ice.

iCLIP2 and iiCLIP were performed by lysing each cell pellet with 1 ml of iCLIP lysis buffer (shared step across all protocols) with the following RNase I dilutions for 3 minutes at 37°C : 0.1U for 1 million cells. For each immunoprecipitation, 2ug of mouse monoclonal antibody (sc-56701) was used with 100ul of protein G beads for 1ml of protein lysate.

### iCLIP2 experiment

iCLIP2 was performed as described in (Buchbender et al. 2019), with the shared crosslinked cells, and RNA fragmentation and immunoprecipitation conditions specified above. Libraries were sequenced on Miseq for SR150 (Table S1)

### iiCLIP protocol

A step-by-step protocol is available upon request by contacting.

Immunoprecipitation and wash conditions are the same as previous iCLIP, as described in (Huppertz et al. 2014). For the specific conditions used in iiCLIP_nd and iiCLIP_d, refer to Table S2.

#### SeqRv adapter ligation

After IP, beads were washed twice in high salt buffer and twice in PNK wash buffer, beads were incubated at 37°C for 40 minutes for 3’end dephosphorylation in the following reaction mix: 8μl 5x PNK pH 6.5 buffer, 1μl PNK (NEB M0201L), 0.5μl FastAP alkaline phosphatase (ThermoFisher, EF0654), 0.5μl RNasin, 30μl nuclease-free water. Beads were washed in PNK wash buffer, and adapter ligation was performed on beads for overnight at 16°C or 75 minutes at 25°C, with the following modified conditions: 6.3μl water, 3μl 10X ligation buffer (no DTT), 0.8μl 100% DMSO, 2.5μl T4 RNA ligase I(M0437M NEB), 0.4μl RNasin, 0.5μl PNK, 2.5μl pre-adenylated adaptor L3-IR-App or L3-XXX-App (stock 1μM), 9μl 50% PEG8000.

#### RecJ adapter removal

Beads were washed twice in high salt buffer and twice in PNK wash buffer after adapter ligation. The beads were then resuspended in the following reaction for adapter removal: 12μl Nuclease-free water, 2μl NEB Buffer 2, 0.5μl 5’ Deadenylase (NEB M0331S), 1μl RecJ endonuclease (NEB M0264S), 0.5μl RNasin, 4μl 50% PEG8000. The beads were incubated at 30°C for 1 hour, then 30 minutes at 37°C for 30 minutes.

#### iCLIP SDS-PAGE and visualisation

Beads were washed twice in high salt buffer and twice in PNK wash buffer after RecJ adapter removal. After washing, beads were then resuspended in 20μl of 1x NuPAGE loading buffer, and the RBP-RNA-linker complexes were eluted at 70°C for 1 minute. This is loaded in a NuPAGE 4-12% Bis-Tris SDS-PAGE, run for 65 minutes at 180V with MOPS buffer, and transferred to a nitrocellulose membrane for 2 hours at room temperature, 30V with 1x NuPAGE Transfer buffer with 10% methanol. The membrane is then visualised at the LI-COR Odyssey-Clx. The pre-stained protein ladder is imaged at the 700nm channel whereas the RBP complexes ligated to the linker is visualised at the 800nm channel (Zarnegar et al. 2016).

#### RNA extraction

The nitrocellulose membrane strips were digested with 10μl proteinase K (Roche, Ref no.: 03115828001) in 200μl proteinase K SDS buffer (10mM Tris-HCl pH 7.4, 100mM NaCl, 1mM EDTA, 0.2% SDS) for 1 hour at 50oC. The supernatant was transferred to a phase lock gel heavy tube (VWR, 713-2536). The RNA extracted were then phenol/chloroform extracted from the supernatant by adding 200μl of Phenol:Chloroform:Isoamyl Alcohol 25:24:1 (Sigma, P3803), incubating shaking for 5 min at 30°C, and centrifuged to separate the aqueous and organic phases. The aqueous phase was further extracted with 0.8ml of chloroform, centrifuged, and the upper phase containing the RNA is transferred to a new tube, and then precipitated with 20μl 3M NaAc, 0.75μl glycoblue and 0.5ml EtOH overnight at −20°C (with or without addition of 0.5μl 1μM barcoded RT primers, Table S1). The RNA is pelleted for 30 minutes at 4°C at 13000G.

#### Reverse transcription and RT primer removal

The RNA is resuspended in 5.5μl (SSIV reaction without PEG8000) or 5μl (for reaction with PEG8000) water then reverse transcribed with barcoded RT primers using the Superscript IV (SSIV) Kit with the following condition:

Add 0.5μl 1μM RT primer, 0.5μl 10mM dNTP, denature at 65°C for 5 minutes, ramp down to 25. Hold at 25°C until addition of 2μl 5x SSIV buffer, 0.5μl 0.1M DTT, 0.25μl RNasin, 0.25μl SSIV, and (see Table S1) 1μl 50% PEG8000. Then incubate in a thermocycler with the following program: 25°C, 5 minutes. 50°C, 5 minutes. 55°C, 5 minutes. 4°C, Hold.

After RT, the reaction incubated with Exonuclease I (2U, add 1μl of a 10-fold dilution of NEB Exonuclease I) for 15 minutes at 37°C. 1.5μl 0.5M EDTA is added to inactivate ExoI, then RNA is degraded by alkaline hydrolysis at 85°C for 15 minutes by adding 1.25μl 1M NaOH. Afterwards, this is neutralised with 1.25μl 1M HCl.

#### cDNA purification with AmPURE beads

cDNA cleanup was performed by adding 45μl (3x) of Agencourt AMPure XP beads and 25μl (1.66x) Isopropanol for capture for 5 minutes at room temperature, then magnetically separated. Supernatant was removed and beads were washed twice, each lasting 30 seconds, by overlaying the beads with 200μl 85% EtOH. After removal and evaporation of EtOH, cDNA was eluted back in 9μl (followed by commercial circligase) or 6.75μl (followed by in-house TS2126) water.

#### cDNA circularisation

Eluted cDNA was moved to a new PCR tube and mixed with CircLigase II (Epicentre) reaction mix containing: 1.5μl 10x CircLigase II Buffer, 0.75μl CircLigase II, 0.75μl 50 mM MnCl2, 3μl 5M betaine, or mixed with in-house TS2126 reaction mix containing: 3μl 5x TS2126 buffer, 0.75μl 50 mM MnCl2, 0.75μl 1mM ATP, 0.75μl purified TS2126, 3μl 5M betaine. Circularisation reaction was incubated at 60°C for 2 hours. Circularised cDNA is purified using the same AMPure XP beads procedure as before. cDNA is eluted in 10μl of water and can be stored at −20°C.

#### iiCLIP input experiment

50μl of input crosslinked lysate was mixed with 50μl of SP3 beads, which were pre-washed and resuspended in water (Moggridge et al. 2018). 100μl of 100% EtOH was mixed with the bead/lysate mixture and incubated for the duration of immunoprecipitation at 4oC for capture of protein-RNA complexes. The input sample then underwent the same iiCLIP procedure as the IP samples beginning from the wash steps before 3’end dephosphorylation. To assay the global input back in the iiCLIP experiment, RNA fragments were extracted from 40-250 kDa on the SDS-PAGE.

### CLIP analysis

All CLIP sequencing data were uploaded to the iMaps webserver (imaps.genialis.com) for standardised primary analysis, for demultiplexing by experimental barcodes, UMI identification, adapter trimming, STAR pre-mapping to rRNAs and tRNAs, STAR alignment to genome, crosslink site assignment using read starts and duplicate removal by UMI sequence. Figures were produced in Rstudio and code is available at (https://github.com/ulelab/iiclip-benchmarking).

### Public eCLIP and iCLIP PTBP1 data

To compare iiCLIP libraries to public data, 2x K562 eCLIP replicates and 4x HeLa iCLIP replicates were merged using bedtools. Fastqs for eCLIP experiments (ENCFF689XJE, ENCFF273VIJ, ENCFF960RXN) were obtained from the ENCODE consortium project (https://www.encodeproject.org/).

### eCLIP pre-processing

The reverse read (R2) was used for analysis as 5’ end of R2 contains the UMI sequence followed by the site at which reverse transcription has truncated. To control for double adapter ligation events, eCLIP R2 fastq files were pre-processed according to the eCLIP SOP (Van Nostrand et al. 2016), with two rounds of cutadapt (performed on iMaps) equivalent to the following parameters:

1st round

-a AACTTGTAGATCGGA -a AGGACCAAGATCGGA -a ACTTGTAGATCGGAA -a GGACCAAGATCGGAA -a CTTGTAGATCGGAAG -a GACCAAGATCGGAAG -a TTGTAGATCGGAAGA -a ACCAAGATCGGAAGA -a TGTAGATCGGAAGAG -a CCAAGATCGGAAGAG -a GTAGATCGGAAGAGC -a CAAGATCGGAAGAGC -a TAGATCGGAAGAGCG -a AAGATCGGAAGAGCG -a AGATCGGAAGAGCGT -a GATCGGAAGAGCGTC -a ATCGGAAGAGCGTCG -a TCGGAAGAGCGTCGT -a CGGAAGAGCGTCGTG -a GGAAGAGCGTCGTGT -m 18 --times 1 --match-read-wildcards --error-rate 0.1 -O 1

2nd round

-a AACTTGTAGATCGGA -a AGGACCAAGATCGGA -a ACTTGTAGATCGGAA -a GGACCAAGATCGGAA -a CTTGTAGATCGGAAG -a GACCAAGATCGGAAG -a TTGTAGATCGGAAGA -a ACCAAGATCGGAAGA -a TGTAGATCGGAAGAG -a CCAAGATCGGAAGAG -a GTAGATCGGAAGAGC -a CAAGATCGGAAGAGC -a TAGATCGGAAGAGCG -a AAGATCGGAAGAGCG -a AGATCGGAAGAGCGT -a GATCGGAAGAGCGTC -a ATCGGAAGAGCGTCG -a TCGGAAGAGCGTCGT -a CGGAAGAGCGTCGTG -a GGAAGAGCGTCGTGT -m 18 --times 1 --match-read-wildcards --error-rate 0.1 -O 5

Pre-processed fastq files were analysed by the primary analysis pipeline on iMaps without demultiplexing and adapter trimming.

### PTBP1 motif coverage

PTBP1 enriched pentamers were previously identified in (Haberman et al. 2017). The top 12 pentamers (“TCTTT”, “CTTTC”, “TCTTC”, “CTTCT”, “TCTCT”, “CTCTC”, “TTTCT”, “TTCTC”, “TTCTT”, “TTTTC”, “TCCTT”, “CTCTT”) were selected for further analysis. Crosslinks weighted by cDNA count with score capping at the 95th percentile on a regional basis were subsampled to ten thousand (five thousand for CDS as this region contains less than ten thousand crosslinks in some libraries) without replacement for calculation of motif coverage around crosslink positions.

### PTBP1 RNA splicing map analysis

Microarray data of PTBP1 KD was taken from (Llorian et al. 2010). Exons were defined using ASPIRE3 comparing KD to WT data to be control exons (|dIRank| < 0.1), silenced by PTB (dIRank < 0.1) or activated by PTB (dIRank > - 0.1). K562 RNAseq data of PTBP1 was taken from ENCODE PTBP1 CRISPR RNA-seq (ENCSR415DJT) and control (ENCSR163JUC). HEK293 RNAseq data of PTBP1/2KD was obtained from a previous publication (Gueroussov et al. 2015). Skipped exons were detected using rMATS (Shen et al. 2014) using only junction counts for K562 and exon and junction counts for HEK293 with a p-value threshold of 0.05 and FDR threshold of 0.1. Silenced and enhanced exons were defined using an inclusion level difference threshold of 0.05; control exons were selected as those with a p-value greater than 0.1, an FDR value greater than 0.1, an mean inclusion level of less than 0.9, and an inclusion level difference less than 0.05. Because rMATS evaluates alternative splicing at the junction/event level, it is possible for the same exon to be duplicated within and across groups. Therefore duplicate entries within each group were collapsed; and exons which overlapped both silenced and enhanced were filtered from the analysis, as well as control exons which overlapped silenced or enhanced exons were removed from the control set. To plot the metaprofile, crosslinks weighted by cDNA counts were score-capped at the 99th percentile to minimise the contribution of outliers. To remove positions which are outliers in crosslinking cDNA counts due to the presence of other abundant RNA species (such as snoRNA), crosslinks which map to regions falling outside of ‘CDS’ and ‘intron’ were masked from the analysis, defined in a hierarchically segmented GTF annotation (based on gencode v27 hg38 annotation) supplied in iMaps computed by the iCount package (see https://icount.readthedocs.io/en/latest/_modules/iCount/genomes/segment.html?highlight=icount). Crosslinking coverage around splice sites were obtained with bedtools coverage function (Quinlan and Hall 2010).

### Motif maps

Silenced exons from HEK293 PTBP1/2 KD RNAseq were separated into 3 classes based on crosslink coverage in the -100…0 window (ROI) upstream of the 3’ splice site: 1) relative crosslink coverage iiclip_d1 > iiclip_nd1, 2) relative crosslink coverage iiclip_d1 < iiclip_nd1, 3) no crosslink detected in window. Relative crosslink coverage was calculated for each exon by counts of crosslinks in ROI divided by the sum of crosslinks in ROI across all silenced exons within the experiment. Motif coverage was visualised as a metaprofile and as a heatmap. In the heatmap, exons were ranked based on the total number of nucleotides within the ROI covered by any PTBP1 motifs to aid visualisation of the distribution of motifs.

### CLIP visualisation

CLIP data were visualised in a comparative manner on individual transcripts or known PTBP1 regulated exons using the software *clipplotr* (https://github.com/ulelab/clipplotr), using the gencode v27 hg38 annotation. Library size normalisation or maxpeak normalisation were performed within *clipplotr* as indicated in figure legends.

---

**Figure S1:**
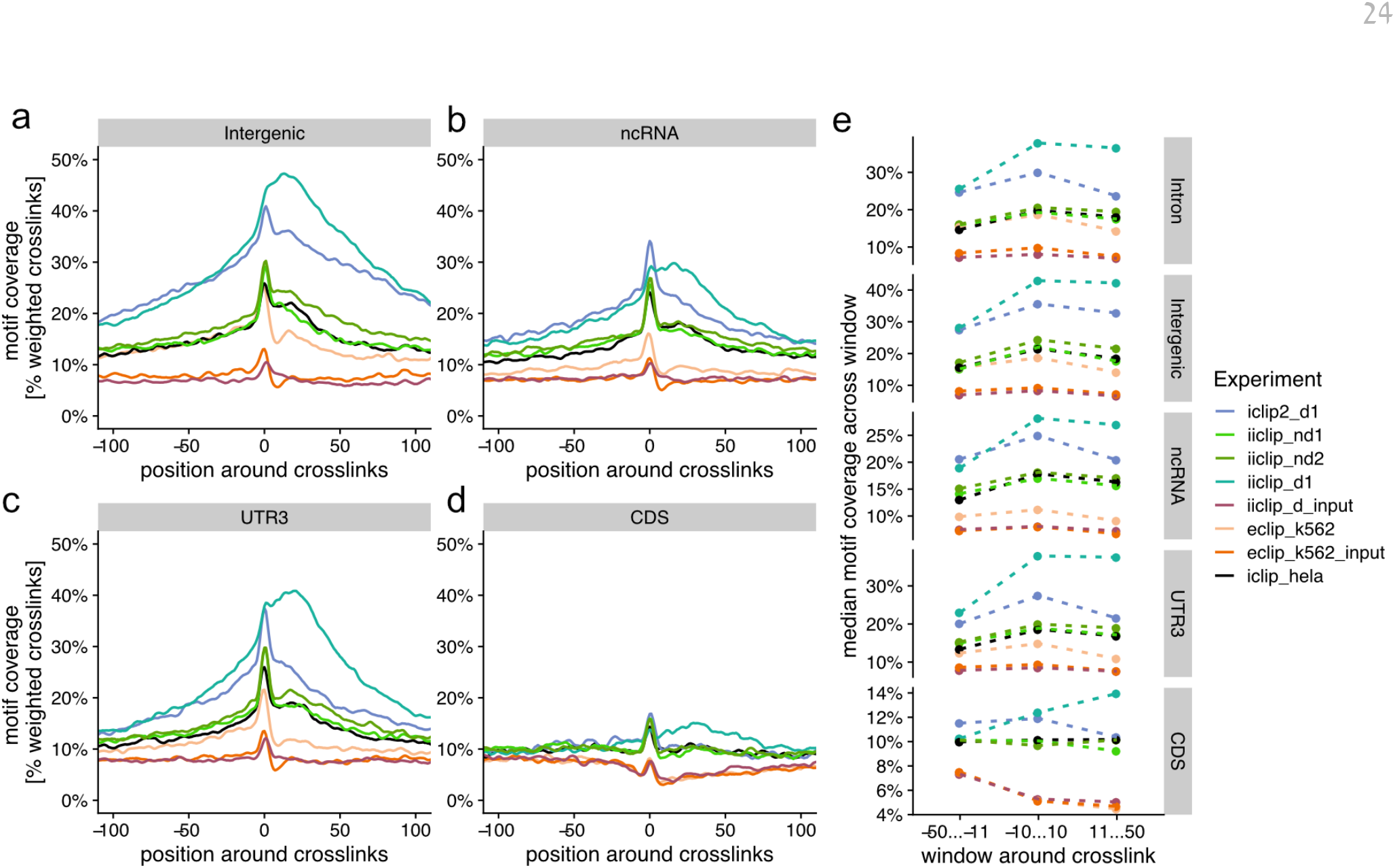
Motif enrichment in PTBP1 and Input crosslink positions across Intergenic, ncRNA, 3’ UTR and CDS regions. Enrichment of known PTBP1 pentamers around weighted crosslink positions in (a) intergenic, (b) ncRNA, (c) 3’ UTR and (d) CDS regions. (e) To summarise the proximal as well as broader enrichment of motif, showing the median motif enrichment of each method in 3 windows upstream, centred or downstream relative to crosslink position.

**Figure S2:**
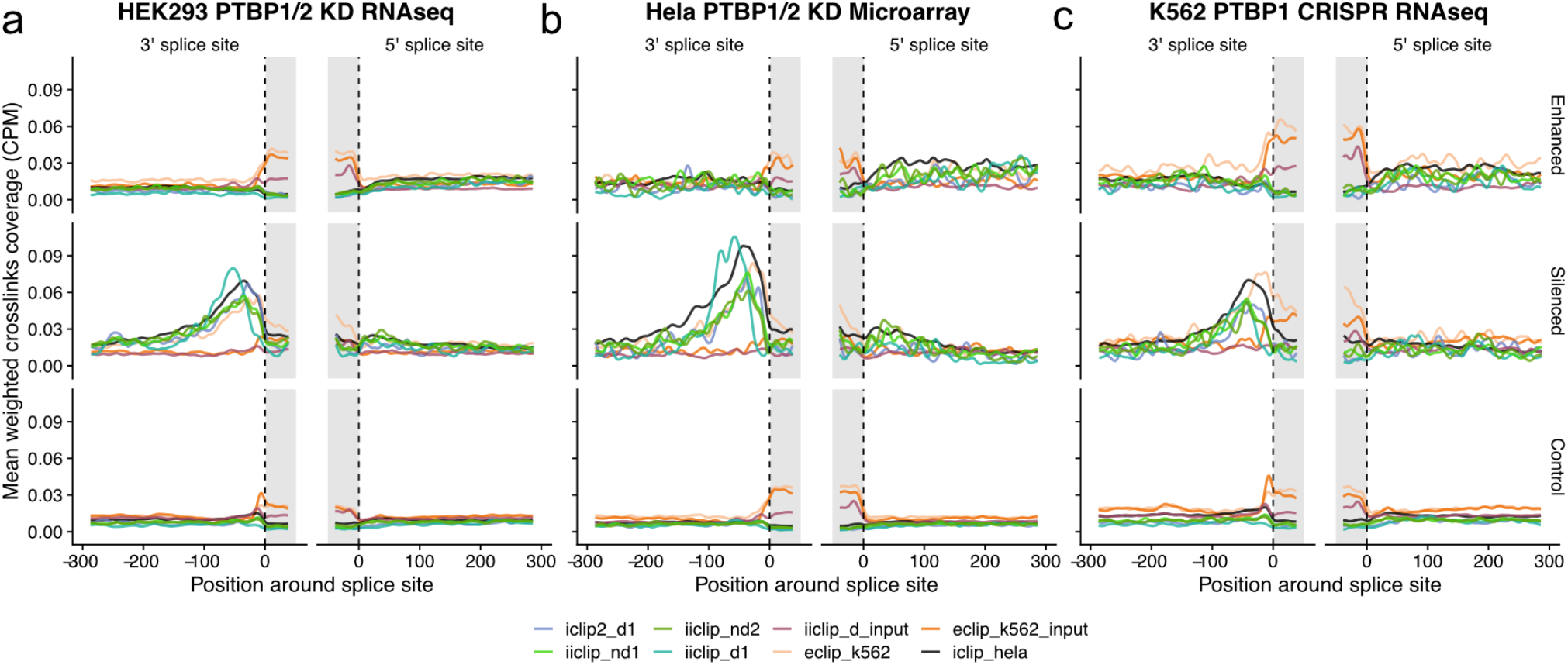
PTBP1 RNA maps evaluate the sensitivity and specificity across methods. PTBP1 RNA maps summarised for enhanced (top), silenced (middle) and control (bottom) exons from 3 public PTB KD datasets: (a) HEK293 PTBP1/2 KD RNAseq analysis (related to Fig.3), (b) HELA PTBP1/2 KD microarray data and (c) K562 PTBP1 CRISPR-treated RNAseq analysis.

**Table S1:** Information on PTB benchmarking experiments.

| Label | Participating lab | Protocol | Replicates | Sequencing | Comments |
| --- | --- | --- | --- | --- | --- |
| iCLIP2 in <u>d</u>eveloper lab (iCLIP2_d) | Konig | iCLIP2 | 1 | SR150, Miseq. | For consistency with other libraries, reads were preprocessed to be truncated to 100 read length |
| iiCLIP in <u>n</u>on-developer lab (iiCLIP_nd) | Turner | iiCLIP | 2 | SR100, Hiseq4000. | without RT primer in ethanol precipitation, RT without PEG8000 and circularisation using commercial circligase II |
| iiCLIP in <u>d</u>eveloper lab (iiCLIP_d) | Ule | iiCLIP | 1 | SR100, Hiseq4000. | with RT primer in ethanol precipitation, RT with PEG8000 and circularisation using in-house TS2126 RNA ligase |

**Table S2:**
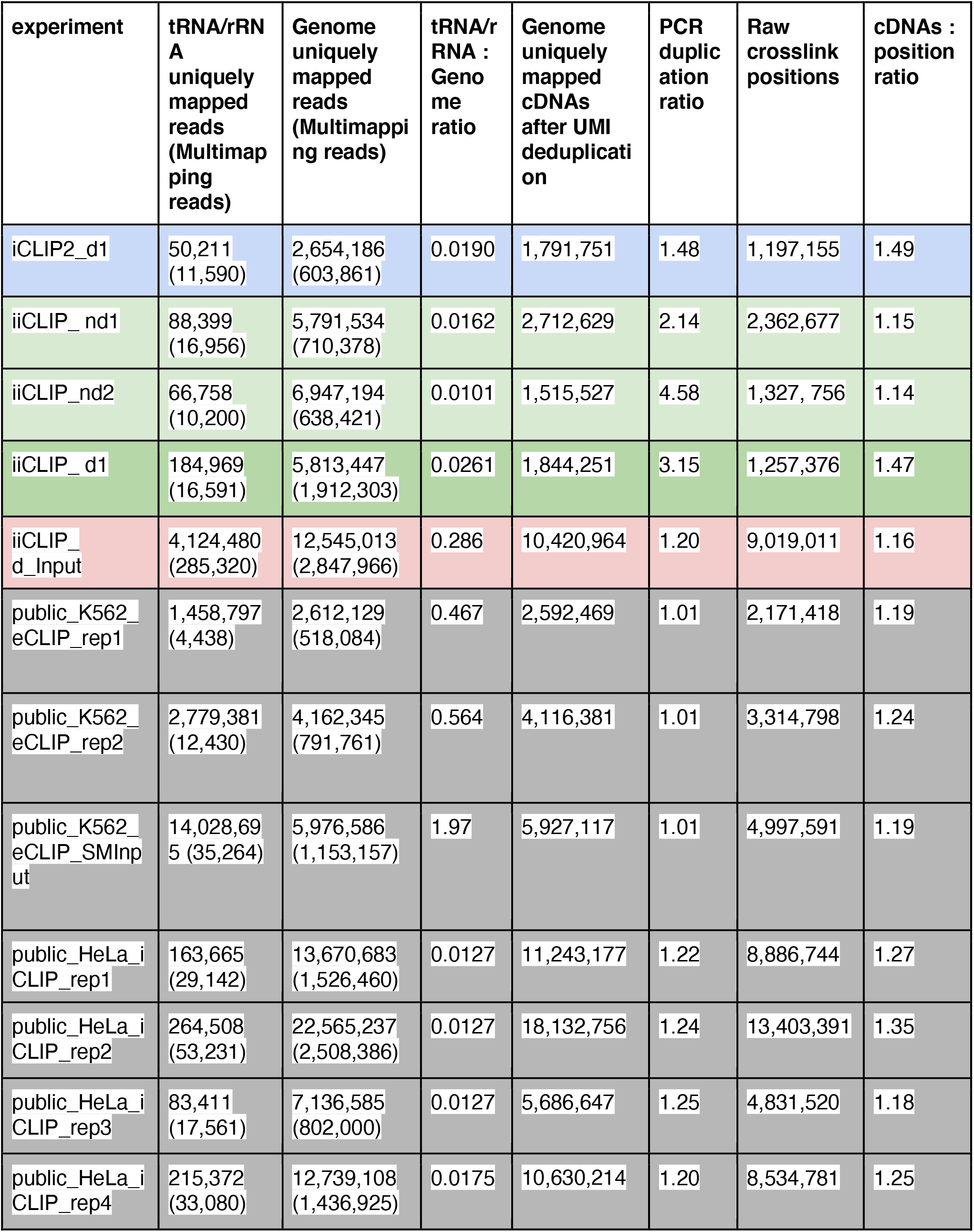
PTBP1 library statistics across community and public datasets.

**Table S3:** source of data.

| experiment | Processed data | Publication |
| --- | --- | --- |
| iCLIP2_d1 | https://imaps.goodwright.com/samples/15590/ | This paper. |
| iiCLIP_nd1 | https://imaps.goodwright.com/samples/15596/ | This paper. |
| iiCLIP_nd2 | https://imaps.goodwright.com/samples/15595/ | This paper. |
| iiCLIP_d1 | https://imaps.goodwright.com/samples/15581/ | This paper. |
| iiCLIP_d_input | https://imaps.goodwright.com/samples/15582/ | This paper. |
| eCLIP_K562 (2 replicates) | https://imaps.goodwright.com/samples/15583/<br>https://imaps.goodwright.com/samples/15584/ | (Van Nostrand et al. 2020) |
| eCLIP_K562_input | https://imaps.goodwright.com/samples/15585/ | (Van Nostrand et al. 2020) |
| iCLIP_hela (4 replicates) | https://imaps.goodwright.com/samples/15586/<br>https://imaps.goodwright.com/samples/15587/<br>https://imaps.goodwright.com/samples/15588/<br>https://imaps.goodwright.com/samples/15589/ | (Attig et al. 2018; Haberman et al. 2017) |

